# Scalable power analysis and effect size exploration of microbiome community differences with Evident

**DOI:** 10.1101/2022.05.19.492684

**Authors:** Gibraan Rahman, Daniel McDonald, Antonio Gonzalez, Yoshiki Vázquez-Baeza, Lingjing Jiang, Climent Casals-Pascual, Shyamal Peddada, Daniel Hakim, Amanda Hazel Dilmore, Brent Nowinski, Rob Knight

## Abstract

Differentiating microbial communities among samples is a major objective in biomedicine. Quantifying the effect size of these differences allows researchers to understand the factors most associated with communities and to optimize the design and clinical resources required to address particular research questions. Here, we present Evident, a package for effect size calculations and power analysis on microbiome data and show that Evident scales to large datasets with numerous metadata covariates.

## Main text

The microbiome has been implicated as a crucial factor in a broad range of health and disease outcomes. Differences in microbial communities have been linked to differential metabolic regulation, often resulting in drastic phenotypic changes. One of the key computational methods for quantifying these community changes is diversity analysis. Alpha diversity measures the overall breadth of microbial features represented in a single sample, while beta diversity quantifies the pairwise community differences between samples via some choice of distance metric. Determining the magnitude of diversity differences among groups of samples is one of the objectives of computational microbiome analysis.

Evaluating the putative differences among groups is most often performed through null hypothesis significance testing (NHST). Under this framework, researchers quantify the probability that an observed difference (or one more extreme) would be observed due to chance (p-value). This value is often used as a measure of significance of diversity differences. However, p-values by themselves are not enough. A significant p-value on its own does not provide any information about the magnitude of a given effect^1^.

In addition to p-values, we suggest reporting the effect size of microbial community differences. Effect sizes quantify differences among sample groups and can be used to describe the magnitude of biological effects. Importantly, standardized effect sizes are dimensionless and can be compared between datasets and experiments^2^. Additionally, effect sizes allow researchers to determine the statistical power of new experimental designs. Statistical power is used to quantify the probability of making a Type II error (false negative) given that the alternative hypothesis is true^3^. Provided an effect size, desired significance level, and sample size, researchers can calculate the statistical power of experimental designs.

Large scale microbiome data collection efforts, such as the American Gut Project (AGP)^4^, TEDDY^5^, and FINRISK^6^ provide a unique opportunity to explore effect sizes across a wide variety of biological factors such as age, obesity, etc. These datasets contain dozens or even hundreds of metadata categories. Determining which covariates contribute the most to microbial diversity is crucial for prioritization of resources. Researchers interested in designing new experiments can keep these effect sizes in mind to efficiently allocate resources to maximize the chances of finding significant biological signal^7^.

Here we introduce Evident, a new open source tool for efficient effect size and power calculations of microbiome data. Evident is available both as a standalone Python package as well as a QIIME 2 plugin^8^. With Evident, researchers can seamlessly explore the effect size of community differences in dozens of metadata columns at once.

Figure 1a shows an overview of the Evident workflow. As input, Evident takes a sample metadata file and a data file. Both univariate data (in vector form such as alpha diversity) and multivariate data (as a distance matrix such as beta diversity) are supported. For univariate measures, the differences in means among groups are considered. For multivariate measures, the difference in means among within-group pairwise distances are considered. It is important to note that Evident does not perform formal hypothesis testing of community differences, only effect size calculations. While we highlight diversity differences in this work, we note that Evident can also be used for other sample-level quantitative metrics such as log-ratios^10^.

**Fig 1:**
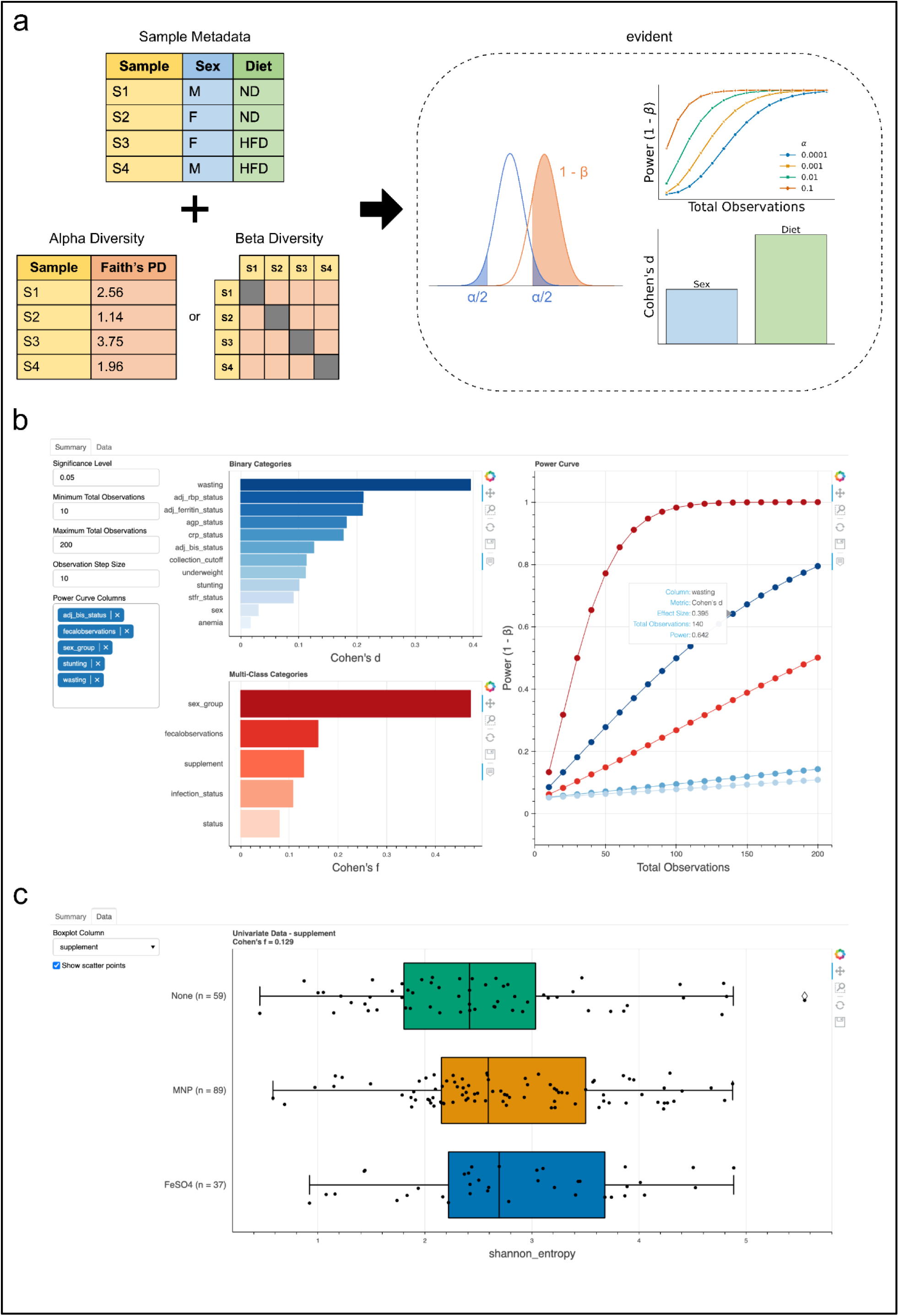
Evident workflow and interactive visualizations. **a**, Graphical overview of Evident usage. Sample metadata with categorical groups are used to determine differences among samples. Effect size calculation can be performed and used to generate power curves at multiple statistical significance levels and sample sizes. **b**,**c** Screenshots of interactive webpage for dynamic exploration of effect sizes and power analysis. Summarized effect sizes of all columns can be used to inform interactive power analysis on multiple groups (b). The underlying grouped data can be visualized with boxplots and, optionally, the raw data as scatter plots (c). Data shown is from McClorry et al. (Qiita study ID: 11402)^9^.

Evident supports both binary categories and multi-class categories. For binary categories, Cohen’s d is calculated between the two levels. For multi-class categories, Cohen’s f is calculated among the levels^11^. Users can specify pairwise effect size calculations between levels of a multi-class category rather than a single group-wise effect size. Effect size calculations can be performed on multiple categories at once with simple parallelization by providing the number of CPUs to use. This architecture allows us to decrease the runtime of effect size calculations for 9495 samples comprising 69 categories from over 12 minutes to 3.5 minutes using 4 CPUs in parallel.

Evident also provides an interactive component by which users can dynamically explore sample groupings. In Figure 1b,c, we show a screenshot of a web app users can access with Evident. Metadata categories are pre-sorted by effect size, allowing efficient determination of interesting categories. Power analysis is implemented dynamically - multiple categories can be visualized simultaneously for a specified significance level and number of observations.

As a demonstration of Evident, we reprocessed 9495 samples from the AGP to compare the published effect sizes with those from a new analysis with Evident^4^. We downloaded the same samples from the original paper and reprocessed the data and metadata in the same manner, focusing on within-group UniFrac^12^ distances. First, we compute the group-wise effect sizes for all valid metadata categories. The top ten binary categories and multi-class effect sizes are shown in Figure 2a,c, respectively. Using these effect sizes, we performed power analyses for each category at a significance level of 0.05 for a range of sample sizes from 20 to 1500 (Figure 2b,d). We plot the distribution of the highest effect size binary and multi-class categories in Figure 2e. Finally, we compute the pairwise effect sizes as performed in the original paper to verify that Evident returns the same values. Figure 2f shows that the effect sizes map extremely closely between the published data and the newly reprocessed data. The values of effect size differences in Figure 2g are distributed around 0, indicating that there is very little difference between effect size calculations.

**Figure 2:**
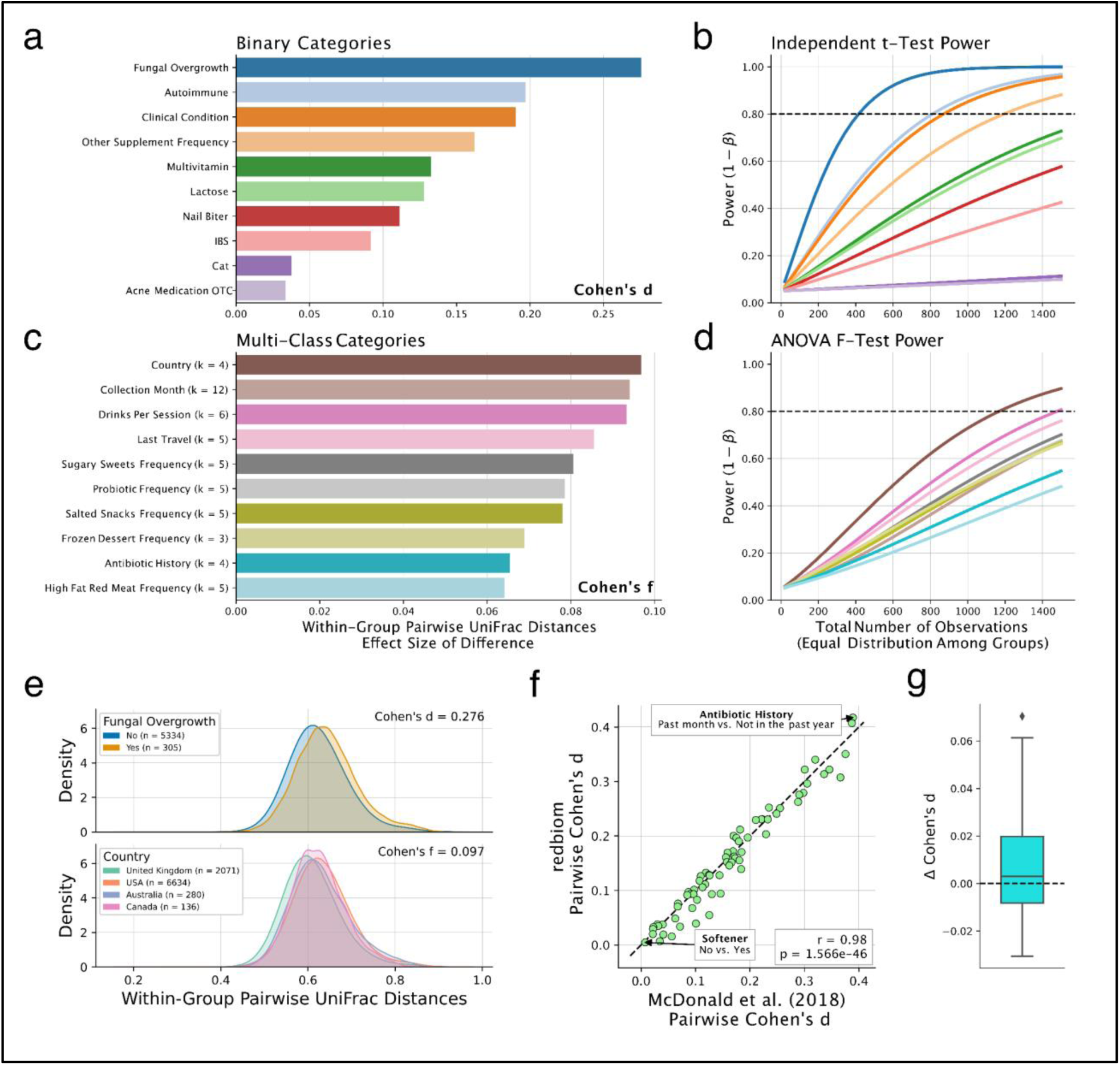
Analysis of American Gut Project data. **a**, Top 10 binary categories by group-wise effect size. **b**, Two-sample independent t-test power analysis of selected binary category effect sizes for significance level of 0.05. **c**, Top 10 multi-class categories by group-wise effect size. **d**, One-way ANOVA F-test power analysis of selected multi-class category effect sizes at significance level of 0.05. **e**, Distributions of within-group pairwise UniFrac distances for highest effect size binary category (top) and multi-class category (bottom). **f**, Comparison of pairwise effect sizes between reprocessed data from redbiom and published effect sizes from McDonald et al. **g**, Boxplot of differences in effect sizes between published and reprocessed effect sizes.

We encourage microbiome researchers to incorporate Evident into their workflows for both reporting effect sizes of microbial community differences and planning experimental designs. In the future, we hope to enhance flexibility by including quantitative metadata categories and variable sample size power analyses.

## Methods

### Overview of Evident

Evident requires a diversity file and its associated sample metadata for a microbiome sequencing experiment. Both univariate and multivariate data are supported as an input pandas Series and scikit-bio DistanceMatrix respectively^13^. When evaluating multivariate differences among sample groups, Evident calculates the difference among pairwise within-group sample distances.

### Effect size calculations

Effect size calculations are available for both binary metadata categories (e.g. Yes vs. No) or multi-class categories (e.g. diet 1, diet 2, diet 3). For binary categories, Evident calculates Cohen’s d; for multi-class categories, Evident calculates Cohen’s f according to a one-way ANOVA. For both types of categories, the pooled standard deviation *(σ* _*p*_) is computed as follows for *G* groups where *n*_*i*_ and *s*_*i*_ are the sample size and sample variance of group *i*, respectively:

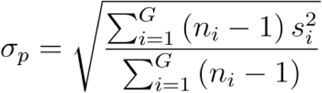

Cohen’s d is calculated by the difference in means of the two groups divided by the pooled standard deviation by the following equation:

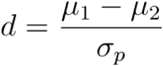

As a rule of thumb, Cohen’s d values of 0.2, 0.5, and 0.8 are generally considered “small”, “medium”, and “large” effect sizes, respectively^11^.

Cohen’s f is calculated by the following equation, where *N* is the total number of samples, and *µ*_*w*_ is the weighted average of all groups by sample size:

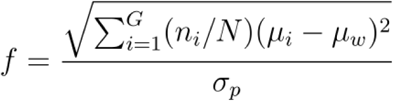

Cohen’s f values of 0.1, 0.25, and 0.4 are generally considered “small”, “medium”, and “large” effect sizes, respectively^11^. For the case of two groups of equal sample size, Cohen’s f is equal to Cohen’s d divided by two.

In Evident, the determination of which effect size measure to use is performed automatically given the number of groups within the chosen metadata column(s). Effect size calculations are performed using NumPy^14^ and SciPy^15^. Calculations in Evident assume that quantitative data is normally distributed and that population variances are homogenous among groups^16^.

Evident allows users to specify the maximum number of levels in a category for the category to be considered. This is useful for ignoring categories with many levels that may not be of interest (e.g. sample identifier). Additionally, one can provide a minimum number of samples in a category level so that rare groups are excluded.

### Power analysis

With computed effect sizes, Evident is able to compute statistical power given significance level(s) and sample size(s). Additionally, Evident allows users to input a target difference (effect size numerator) to use in lieu of automatic computation. This is useful for researchers interested in designing experiments with specific effect sizes in mind^7^. Power analysis assumes that the number of samples is the same in each group for both of these statistical tests. These power analyses are calculated using the statsmodels package in Python^17^.

Notably, Evident is designed for flexibility in power analysis. Users can compute either number of observations, effect size, or statistical power given the other two variables. Additionally, Evident is designed with generating power curves in mind. For example, with a Cohen’s d of 0.4, a user can specify significance levels of 0.1, 0.05, and 0.01 from 20 total observations to 100 in increments of 10. The statistical power will then be evaluated at each entry in the Cartesian product of these two parameter sets. These results can be directly plotted as a power curve using Evident, delineating the curves from different significance levels.

### Repeated measures

Evident supports a limited implementation of repeated measures analysis. For datasets in which the same subject is measured more than once, statistical analysis must be performed with this in mind to account for variation due to between and within subjects. Univariate data such as alpha diversity can be analyzed with repeated measures in mind by providing a mapping of sample to subject.

For repeated measures datasets, Evident calculates eta squared *(η*^2^). This effect size can be used to calculate statistical power of a balanced repeated measures ANOVA. The number of subjects and number of measurements per subject are used to determine the total sample size. In addition to significance level, sphericity (variance between pairs of treatments^18^) and sample correlation parameters are used to calculate statistical power as described previously^19,20^. For convenience, subjects with missing values are removed.

### Interactive exploration of community differences

The interactive visualization provided in Evident is created with Bokeh. Given microbiome data and sample metadata, Evident creates a Bokeh app that dynamically calculates effect sizes and power analysis for the chosen parameters. This view also shows the raw data values as boxplots with optional scatter points.

### Analysis of AGP data

A sample ID list was generated from the original distance matrix used in the AGP study. 100 nucleotide 16S-V4 data for these samples were downloaded from the AGP study on Qiita (study ID: 10317) using redbiom^21,22^. Both preparation and sample metadata were also retrieved with redbiom. Due to multiple preparations of some samples, we performed disambiguation by keeping the samples with the highest sequencing depth.

We then processed the feature table and metadata according to the original study. The original workflow used the default parameters in Deblur to remove features with fewer than 10 occurrences in the data^23^. Because Qiita does not perform this filtering by default, we performed this filtering manually. To remove sequences associated with sample bloom, we performed bloom filtering^24^. We then rarefied the feature table to 1250 sequences as in the original analysis.

We processed the sample metadata in accordance with the original study. Because of differences in self-reporting protocols from 2018, metadata categories associated with reported Vioscreen responses as well as those associated with alcohol consumption were removed. The following categories were removed due to mismatches in sample metadata: roommates, allergies, age_cat, bmi_cat, longitude, latitude, elevation, height_cm, collection_time, and center_project_name. Only the top four annotated countries were considered - US, UK, Australia and Canada. All other countries were ignored. Overall, 69 metadata categories common to both the original data and redbiom data were used for further analysis.

Sequences from the feature table were placed into a 99% Greengenes^25^ insertion reference tree using SEPP^26^. We then used unweighted UniFrac to generate a sample-by-sample distance matrix^27^. This distance matrix was used as input to Evident along with the disambiguated, processed sample metadata.

We used effect_size_by_category to calculate the whole-group effect sizes for each column in the metadata and pairwise_effect_size_by_category to calculate the group-pairwise effect sizes for multi-class categories. For each whole-group effect size, we computed a power analysis for alpha values of 0.01, 0.05, and 0.1. Power was calculated on total sample size values from 20 to 1500 in increments of 40 samples. Evident analyses were performed in parallel on a high performance computing environment. Group-wise and pairwise effect size calculations both took under 4 minutes for 94 metadata categories on 9495 samples using 4 CPUs (we note the AGP paper used n=9511 but operated at 125nt; we observe a slightly reduced number of samples at 100nt). We also benchmarked group-wise effect size calculations using only a single CPU as comparison; this process took 12.4 minutes - meaning the parallelization decreased runtime by approximately 3.5x. Power analysis calculation took 2 minutes for 94 categories using 8 CPUs in parallel.

## Code availability

The latest version of Evident is available at https://github.com/biocore/evident under the BSD-3 license. Evident is installable from PyPI both as a standalone Python 3 package and a QIIME 2 plugin. The scripts used to download and analyze AGP data as well as the processed Evident results are available at https://github.com/knightlab-analyses/evident-analyses. Analysis of AGP data in this study was performed with Evident version 0.2.0.

## Data availability

Data for the demonstration in Figure 1 were downloaded from Qiita (study ID: 11402)^9^ at 90 nucleotides using the deblur^23^ pipeline. AGP data were downloaded from Qiita (study ID: 10317) using redbiom with context “Deblur_2021.09-Illumina-16S-V4-100nt-50b3a2”. The original pairwise effect sizes, sample metadata, and unweighted UniFrac distance matrix were downloaded from the original McDonald et al. study for comparison.

## Contributions

G.R., A.G, D.M., & R.K. conceived the idea for the software and study. G.R., A.G., D.M., Y.V.B & L.J. developed the software. G.R., D.M., & A.G. conducted the analysis of the AGP data.

B.N. assisted with AGP metadata curation. A.H.D., Y.V.B., D.M., & D.H. reviewed the software code and provided valuable feedback and bug reports. C.C., L.J., & S.P. contributed to the statistical computation used in Evident. All authors contributed to and reviewed the final manuscript.

## Acknowledgements

We would like to thank the members of the Knight Lab for feedback on the scope and details of Evident. We thank Jamie Morton for valuable discussions about effect size. This work was supported in part by the Alfred P. Sloan foundation (G-2017-9838), NIH-NIDDK (P01DK078669), NIH-NCI (U24CA248454), and NIH (1DP1AT010885, U19AG063744).

Research of S.P. was funded by the intramural research program of *Eunice Kennedy Shriver* National Institute of Child Health and Human Development, NIH.

